# Laminar microcircuitry of visual cortex producing attention-associated electric fields

**DOI:** 10.1101/2021.09.09.459639

**Authors:** Jacob A. Westerberg, Michelle S. Schall, Alexander Maier, Geoffrey F. Woodman, Jeffrey D. Schall

**Affiliations:** Department of Psychology, Center for Integrative and Cognitive Neuroscience, Vanderbilt Vision Research Center, Vanderbilt Brain Institute, Vanderbilt University, Nashville, Tennessee 37240, USA; Centre for Vision Research, Vision: Science to Applications Program, Departments of Biology and Psychology, York University, Toronto, Ontario M3J 1P3, Canada

**Keywords:** CSD, ECoG, EEG, LFP, N2pc, V4

## Abstract

Cognitive operations are widely studied by measuring electric fields through EEG and ECoG. However, despite their widespread use, the component neural circuitry giving rise to these signals remains unknown. Specifically, the functional architecture of cortical columns which results in attention-associated electric fields has not been explored. Here we detail the laminar cortical circuitry underlying an attention-associated electric field often measured over posterior regions of the brain in humans and monkeys. First, we identified visual cortical area V4 as one plausible contributor to this attention-associated electric field through inverse modeling of cranial EEG in macaque monkeys performing a visual attention task. Next, we performed laminar neurophysiological recordings on the prelunate gyrus and identified the electric-field-producing dipoles as synaptic activity in distinct cortical layers of area V4. Specifically, activation in the extragranular layers of cortex resulted in the generation of the attention-associated dipole. Feature selectivity of a given cortical column determined the overall contribution to this electric field. Columns selective for the attended feature contributed more to the electric field than columns selective for a different feature. Lastly, the laminar profile of synaptic activity generated by V4 was sufficient to produce an attention-associated signal measurable outside of the column. These findings suggest that the top-down recipient cortical layers produce an attention-associated electric field capable of being measured extracranially and the relative contribution of each column depends upon the underlying functional architecture.

## Introduction

Research into extracranial electric fields provides fundamental insights into the mechanisms of human perception, cognition, and intention. For instance, event-related potential (ERP) components like the N2pc (Luck and Hillyard, 1994; Eimer, 1996; Woodman and Luck, 1999) and Pd (Hickey et al., 2009) reliably index selective attention in humans and monkeys, alike. However, the interpretation of these extracranial measures of attention is severely limited by uncertainty about the exact neural processes that generate these signals (Nunez and Srinivasan, 2006). Understanding what brain processes an electric field indicates requires knowing how it is generated (e.g., Cohen, 2017).

One avenue to localize neural generators of electric fields is through inverse source localization (Michel et al., 2004; Grech et al., 2008). However, the results are indefinite and cannot offer conclusive answers. Moreover, these methods do not allow for the probing of the underlying neural circuitry. For example, most EEG signals are hypothesized to be generated by interlaminar interactions in cortical columns (Nunez and Srinivasan, 2006). Columnar microcircuits are ubiquitous across the brain (Douglas et al., 1989; Douglas et al., 1991 but see Godlove et al., 2014), having a well-defined anatomical structure (Mountcastle, 1997; Kaas, 2012) and consistent physiological activation pattern (Bastos et al., 2012). This canonical cortical microcircuitry allows for a framework in which to interpret columnar dynamics in sensory or cognitive tasks, yet the relationship between this functional architecture and electric fields related to cognition commonly measured in humans has yet to be explored.

Electric fields measured at the surface of the brain (ECoG) and scalp (EEG) are theorized to stem from dipoles in cortex. However, measuring current dipoles requires sampling electrical potentials across all the layers of the cerebral cortex. Such laminar neurophysiological measurements are rare and unsystematic in humans. Work in rodents has uncovered intriguing insights into cortical laminar microcircuits underlying evoked EEG signals, but all of these were limited to sensory responses (Jellema et al., 2004; Bruyns-Haylett et al., 2017; Næss et al., 2021). Fortunately, macaque monkeys produce homologues of the attention-associated EEG signals (N2pc: Woodman et al. 2007; Cohen et al., 2009; Purcell et al., 2013; Pd: Cosman et al., 2018). Laminar neurophysiological measurements (Schroeder et al., 1998; Maier et al., 2010; Buffalo et al., 2011; Hansen et al., 2011; Self et al., 2013; Godlove et al., 2014; Engel et al., 2016; Klein et al., 2016; Hembrook-Short et al., 2017; Nandy et al., 2017; Trautmann et al., 2019; Westerberg et al., 2019; Tovar et al., 2020; Ferro et al., 2021) and EEG (Schmid et al., 2006; Woodman et al., 2007; Sandhaeger et al., 2019) are well established in macaques. However, despite many studies linking intra- and extracranial signals (Schroeder and Givre, 1992; Whittingstall and Logothetis, 2009; Musall et al., 2014; Snyder and Smith, 2015), to date, little is known about the laminar origins of ERPs in primates.

Here we show that visual cortex generates dipoles through layer-specific transsynaptic currents that give rise to electric fields that track the deployment of selective attention. These dipoles were generated by the extragranular compartments of cortex – indicating these cognitive operations likely arise from top-down interactions. Moreover, functional architecture – in the form of feature columns – were associated with the relative contribution of individual, local cortical columns to the global electric field. These results are the first to our knowledge to describe laminar specificity in synaptic activations contributing to the generation of electric fields associated with cognitive processing.

## Results

### Attention task

To investigate extracranial manifestations of attention-associated electric fields, we first trained macaque monkeys to perform a visual search task (Figure 1A). Three macaque monkeys (designated Ca, He, and Z) performed visual search for an oddball color target (red or green), presented within an array of 5 or 7 uniform distractors (green or red) (N sessions: monkeys Ca, 21; He, 9; Z, 18). A fourth monkey (P) performed visual search for an oddball shape (T or L) presented within an array of up to 7 uniform distractors (L or T) (N sessions: monkey P, 22). Each animal performed well above chance [chance level for monkeys Ca, He: 16.6%; P, Z: 12.5%] (behavioral accuracy in color search: monkeys Ca, 88%; He, 81%; Z, 85%; accuracy in shape search: monkey P, 66%). We used a color pop-out search in our initial recordings so that we could be certain of which item received the benefit of attention in the array, and we used the more difficult search data determine the generality of our findings.

**Figure 1.**
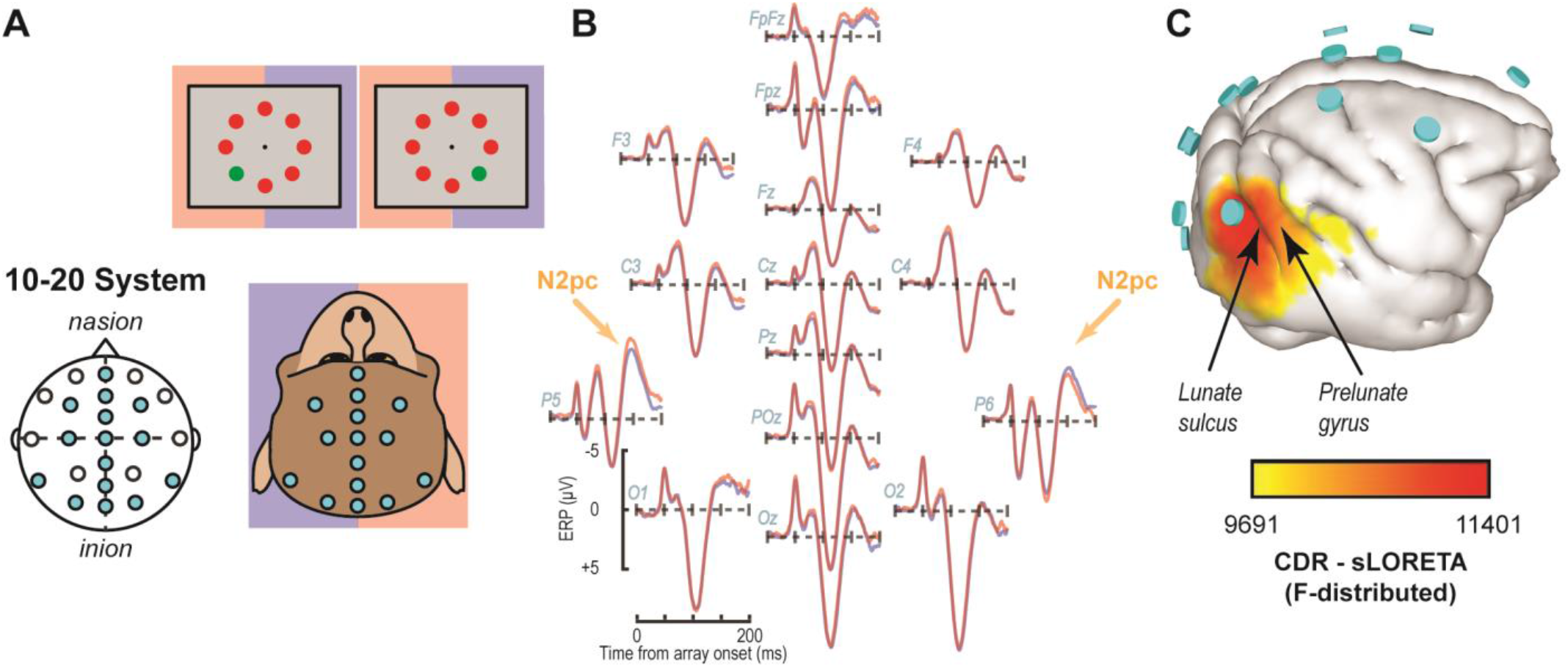
EEG traces and inverse source localization for the N2pc index of attention in monkeys. (A) EEG was recorded from electrodes arranged according to the 10-20 system in monkeys performing visual search for a colored oddball stimulus (see monitor diagrams showing two example search arrays in top panels). Blue and red shading indicates the relationship between visual hemifields and cerebral hemispheres to highlight the mapping between lateralized EEG signals and target location. (B) Trial-averaged EEG traces from monkey Z following presentation of search arrays with the target in either the right (blue) or left (red) visual hemifield. The voltage differential that characterizes the N2pc arises ∼150 ms after array presentation for the target in the left vs. right visual fields (orange arrows). The N2pc was significant at posterior sites P5 and P6 (dependent samples t test between response polarizations averaged between 125-250 ms following array onset for contralateral and ipsilateral target presentations (t(35) = 2.42, p = 0.02)). (C) Inverse solution using sLORETA to determine current distribution consistent with voltage distribution during the N2pc (113-182 ms) when the target was in the left hemifield. Current density is displayed over the 3D boundary element model derived from a magnetic resonance scan of monkey Z. Data was clipped below the 85% maximum value for display purposes. Cyan disks indicate EEG electrode positions. Current density is concentrated beneath electrode P6 caudal to the lunate sulcus and in area V4 on the prelunate gyrus. Both results in B and C are reproduced for a second monkey in Figure S1.

### Inverse modeling of attention-associated extracranial electric fields points to visual cortex

Once animals could successfully perform visual search, we implanted an array of electrodes approximating the human 10-20 system in monkeys P and Z (Figure 1A). Using these electrodes, we observed extracranial electric dynamics in both monkeys. An index of attention known as the N2pc manifests during visual search. The N2pc served as our representative attention-associated electric field indicating attention in this task. The magnitude of the N2pc was largest over occipital sites (Figure 1B, S1), consistent with previous reports in humans and macaques (Luck and Hillyard, 1994; Eimer, 1996; Woodman and Luck, 1999; Hopf et al., 2000; Woodman et al., 2007; Cohen et al., 2009; Purcell et al., 2013). Next, we used sLORETA inverse modeling for source localization. Previous source estimates for the N2pc identified the human homologue of V4 (Luck and Hillyard, 1990, 1994; Hopf et al., 2000). These findings are consistent with numerous reports that areas in mid-level visual cortex in monkeys produce robust attention signals (Moran and Desimone, 1985; Luck et al., 1997; McAdams and Maunsell, 1999; Reynolds et al., 1999; Fries et al., 2001; see Roe et al., 2012 for review) across cortical layers (Engel et al., 2016; Nandy et al., 2017). In line with these earlier studies, our inverse models showed that current sources include V4 on the prelunate gyrus (Figure 1C, S1). However, the modeled current sources also included other cortical regions, as is common for inverse solutions. Notably, the inverse solution identifies V1 to be about as strong as V4 in contributing to the N2pc, which is unlikely given current knowledge on attentional modulation for each area (Motter, 1993; Luck et al., 1997; Kastner et al., 1999; Buffalo et al., 2010). Given the primary feature used in the search task was color, we decided to investigate the laminar profile of attention-associated electric field generation in V4 where color is better represented (Roe et al., 2012).

### V4’s laminar microcircuit produces dipoles that predict the attention-associated electric field

Guided by magnetic resonance imaging, linear electrode arrays (LMAs) were inserted into area V4 of two monkeys, Ca and He. LMAs were placed perpendicular to the cortical surface, spanning supragranular (L2/3), granular (L4), and infragranular (L5/6) cortical layers. Simultaneously, an extracranial electric signal was recorded immediately above V4 – critically the recording took place outside of the cortical column itself. Current source density (CSD) was derived from the local field potentials (LFPs) sampled across V4 layers. To relate the extracranial signal (Figure 2A) to synaptic currents estimated as CSD (Figure 2B-D), we employed information theory to capture multivariate factors and nonlinearities between signals (Shannon, 1948; Cover and Thomas, 2006). Importantly, information theory analyses are model independent (Timme and Lapish, 2018). Information theory thus is superior to standard linear models since these models cannot capture all potential relationships between signals. The relationship between the extracranial signal and CSD were assessed in four discrete steps, as illustrated by a representative session (Figure 2E-F, S2). Again, we use the time period of the purported N2pc as our primary focus for determining whether V4’s laminar circuitry is involved in the production of attention-associated electric fields.

**Figure 2.**
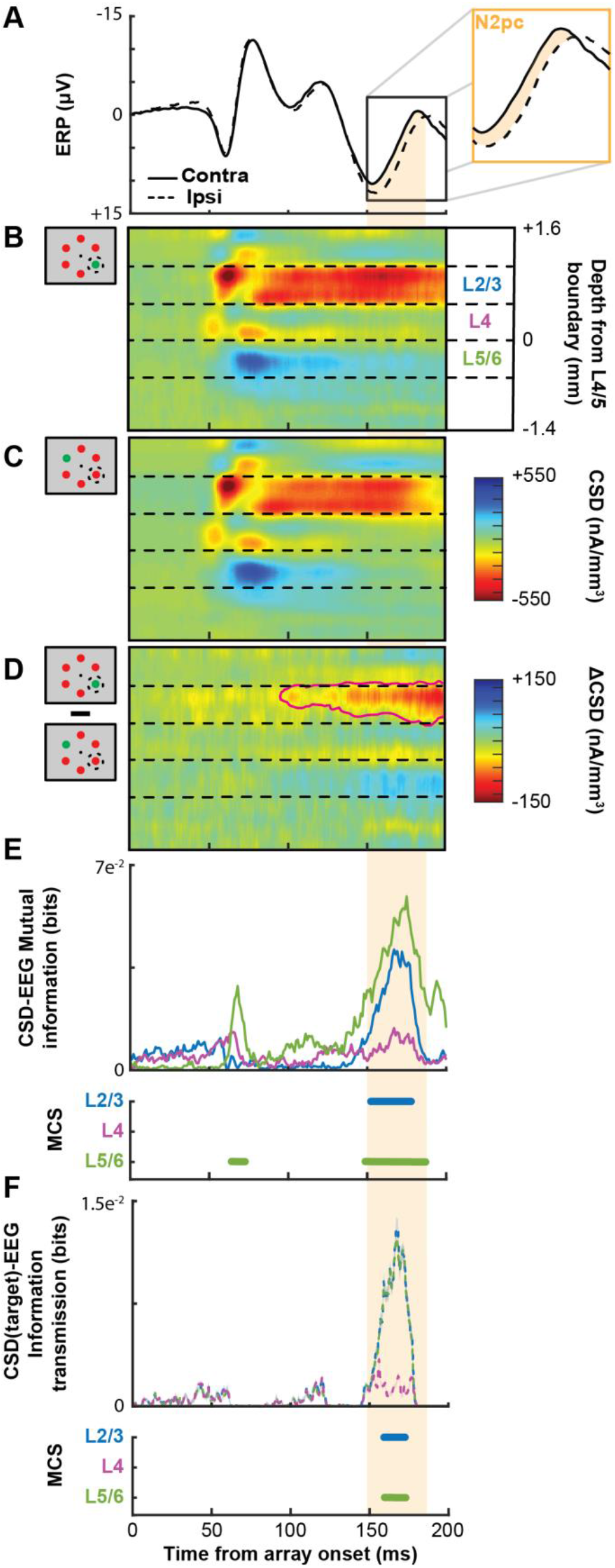
Extracranial attention-associated signal and simultaneously recorded V4 synaptic currents during representative session. (A) Extracranial ERPs as voltages aligned to array onset, averaged over all trials when the target was presented contra-(solid) or ipsilateral (dashed) to the electrode. Inset magnifies the N2pc window, highlighted in orange, defined as the difference in potentials 150–190 ms following array onset. (B) Cortical (laminar) current source density (CSD), aligned on array presentation when the target appeared in the population receptive field of the column. Dashed lines delineate estimated boundaries between supragranular (L2/3), granular (L4), and infragranular (L5/6) layers. CSD values were interpolated and smoothed along depth for display only. Current sinks are indicated by hotter hues and current sources by cooler hues, respectively. The earliest sink arises in putative L4, likely from rapid feedforward transmission. (C) CSD evoked by target outside the receptive field. (D) Subtraction of CSD responses shown in B and C. The only statistically significant differences (determined through a t test across time with p < 0.05, outlined by magenta line) were due to a current sink in L2/3 that arose gradually ∼100 ms after array presentation. This relative sink was associated with a weak relative source in L5/6. (E) Mutual information between CSD and the extracranial signal for L2/3 (blue), L4 (purple), and L5/6 (green), aligned on array onset. Timepoints with significant mutual information were computed through Monte Carlo shuffle simulations (MCS). Epochs with significant mutual information persisting for at least 10 ms are indicated by horizontal bars. No such epochs were observed in L4. Highlighted region indicates period of N2pc. (F) Information transmission about target position from V4 CSD to the extracranial signal. Conventions as in E.

First, we employed Monte Carlo simulations of the mutual information analysis to verify that the extracranial signal exhibits significantly enhanced information about target position during the time window of the N2pc (Figure S2). Second, we measured target information across the layers of V4 during the N2pc temporal window. This analysis revealed enhanced information in L2/3 and L5/6 but not in L4 (Figure S2). Third, we computed the mutual information between the extracranial signal and CSD during the N2pc window, irrespective of target position. This analysis showed a significant relationship between extracranial signal and the CSD in L2/3 and L5/6 but not in L4 (Figure 2E, S2). Fourth, we measured the transmitted information about target location from CSD to extracranial signal during the N2pc window (Timme and Lapish, 2018). This analysis demonstrated significant information transmission to the extracranial signal from L2/3 and L5/6, but not L4 (Figure 2F, S2).

Averaged across sessions, we observed that the attention-associated electric field during the N2pc window (Figure 3A) was associated with a consistent CSD pattern (Figure 3B). This relationship was observable in each monkey (Figure S3A-B). Presentation of the search array in any configuration elicited an early current sink in L4, followed by a prolonged sink in L2/3 that was associated with a briefer source in L5/6.

**Figure 3.**
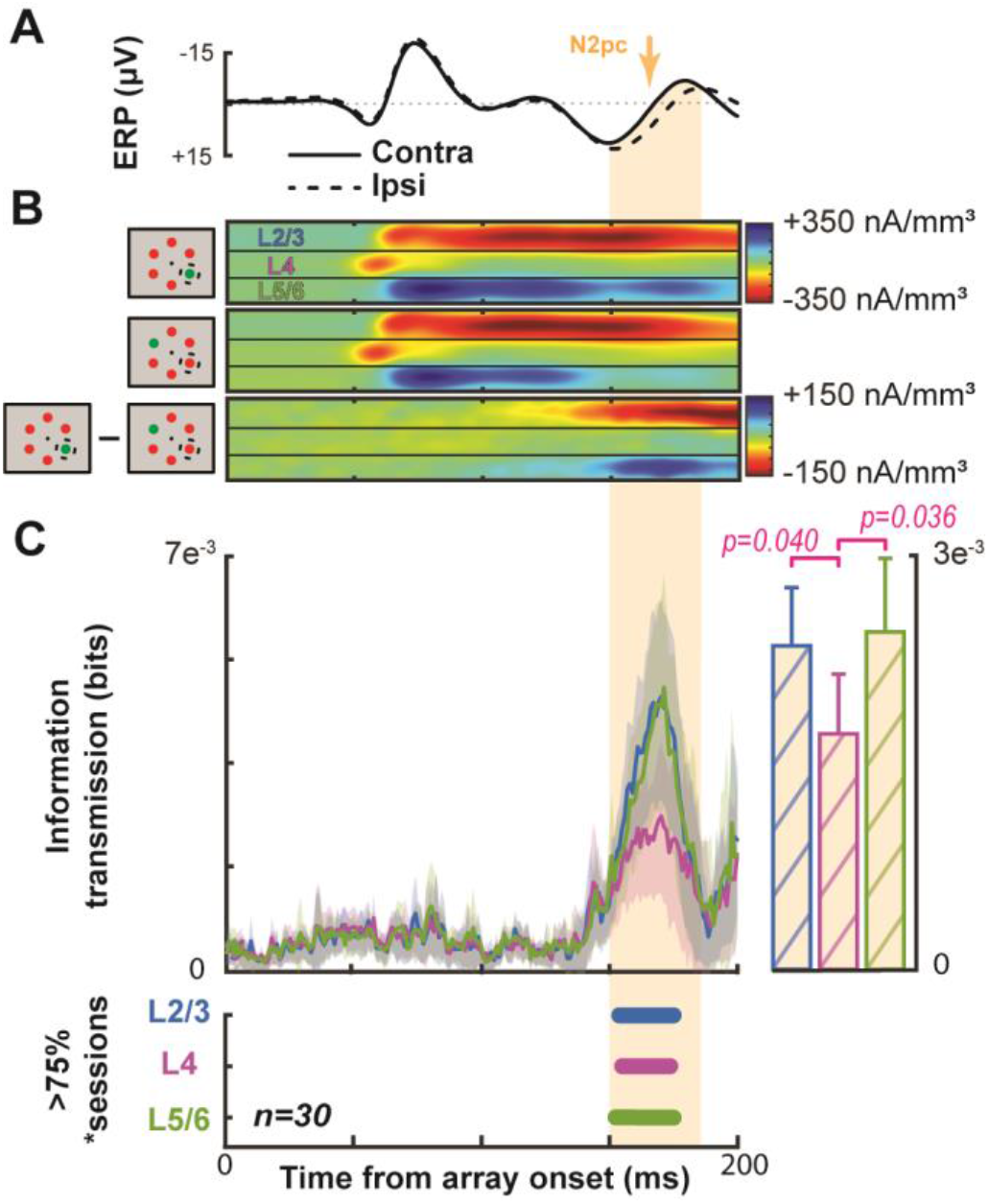
Grand average demonstrating the link between V4 CSD and the extracranial attention-associated electric field. Conventions as in Figure 2. (A) Average ERP across all sessions and animals with the target contra- (solid) or ipsilateral (dashed). The N2pc interval is indicated by orange shading. (B) Average V4 CSD with the target in (top) or out of the RF (center) with the difference between the two at the bottom. (C) Grand average information transmission about target position from V4 layers to the extracranial signal as a function of time (left). Average +2 SEM of information transmission during the N2pc window (right). Panel below shows that Information transmission from L2/3 and in L5/6 was significantly greater than that from L4 (t test p < 0.05). Timepoints with significant information transmission were assessed through Monte Carlo simulations during >75% of sessions. Epochs with significance persisting for at least 10 ms are indicated by horizontal bars, color coded for each laminar compartment (bottom).

We next computed information transmission about target location from the CSD to extracranial signal for each session (Figure 3C). All cortical layers provided significant information transmission in >75% of sessions during the N2pc window (150-190 ms following array onset). However, the magnitude of transmitted target information was significantly greater in L2/3 and L5/6 relative to L4 (L2/3-L4: t(29) = 2.15, p = 0.040; L5/6-L4: t(29) = 2.20, p = 0.036). The magnitude of information transmission was not significantly different between L2/3 and L5/6 (t(29) = 0.21, p = 0.84).

Across sessions, the three other information theoretic analyses were consistent with the example session (Figure S2). Significant information transmission during the time of the N2pc was observed in each monkey (Figure S3C). Thus, the current dipole in V4 generated by the L2/3 CSD sink and the L5/6 CSD source contributes to the N2pc measured in the overlying extracranial electric field.

### Feature selectivity determines columns’ relative roles for attention-associated electric field generation

Given the selectivity of V4 neurons for color (Figure 4A) (Roe et al. 2012) and the homogenous columnar representation of V4 color selectivity (Zeki, 1973, 1980; Tootell et al., 2004; Conway and Tsao, 2009; Kotake et al., 2009), we next investigated the role of columnar color tuning for the contribution of that column to the attention-associated electric field. To quantify color selectivity through depth, we computed the response ratio between red and green stimuli (Figure 4B). Responses were measured as power in the gamma range (30-150 Hz) because this activity reflects local circuit interactions (Ray and Maunsell, 2011) as well as feature selectivity in visual cortex (Berens et al., 2008) and is more reliably measured across all LMA contacts than spiking activity.

**Figure 4.**
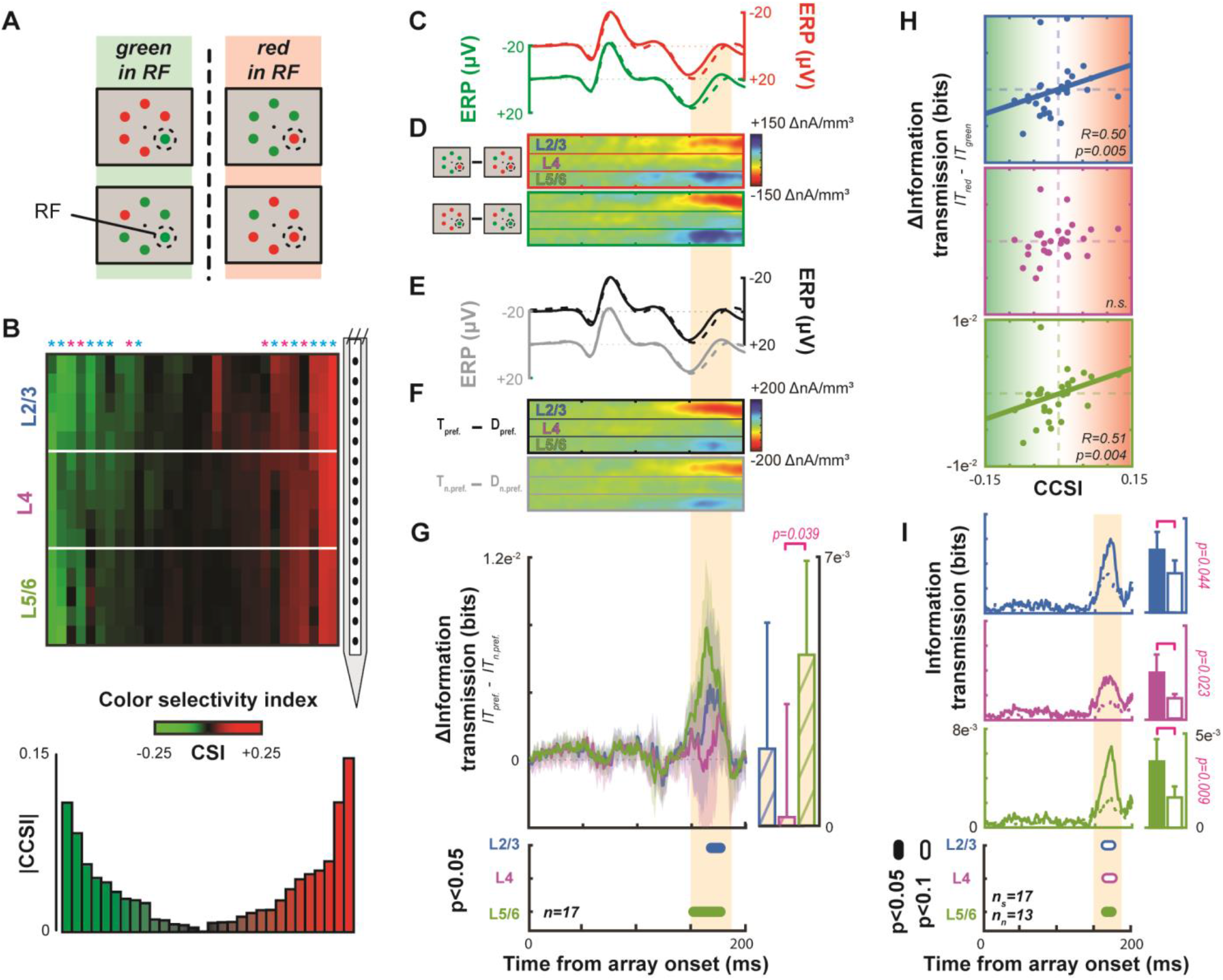
Contribution of columnar feature selectivity to the attention-associated electric field. Conventions as in Figure 2. (A) Visual search array configurations used for color selectivity analyses. (B) Laminar profiles of red/green color selectivity across all sessions session. The hue of each point across cortical depth signifies the value of a color selectivity index (CSI), derived from local gamma power. CSI values <0 indicate preference for green, and values >0, preference for red. CSI is smoothed across 2 adjacent channels for display only. Sessions are sorted from left to right based on a second index that estimates each column’s combined selectivity, termed column color selectivity index (CCSI). A bar plot representing each session’s CCSI is plotted below. Asterisks indicate columns that were significantly color-selective (Wilcoxon sign rank, p < 0.05). Asterisk color indicates which monkey the column was recorded from (monkey Ca: cyan; He: magenta). (C) Average ERPs for trials when a red (top) or green (bottom) target or distractor appeared in the RF based on the 17 significantly color selective columns. Conventions as in Figure 2. (D) Difference in CSD when the target appeared within the columnar population receptive field (RF), compared to out-of-RF trials when a red (top) or green (bottom) target or distractor appeared in the RF (n = 17). (E) Average ERP for trials when the preferred color (top) or non-preferred color (bottom) target or distractor appeared in the RF for the 17 color selective columns. Conventions as in Figure 2. (F) Difference in CSD when the target was within vs. out of the RF, for trials when the preferred color (top) or non-preferred color (bottom) target or distractor appeared in the RF. (G) Average across color-selective columns for subtraction of information transmission from laminar CSD to the extracranial signal about non-preferred color target position from information transmission about preferred color target position. Conventions as before. L2/3 and L5/6 but not L4 contribu te significantly to the extracranial signal. (H) Correlation plots between the CCSI for each session and the difference in information transmission between the red and green stimulus conditions for L2/3 (blue, top), L4 (purple, center), and L5/6 (green, bottom). Spearman correlation reported in lower righthand corner of each plot. Data from all 30 sessions included. (I) Comparison of feature selective (solid line, n _s_ = 17) and non-feature-selective (dashed line, n_n_ = 13) columns for each laminar compartment (L2/3: blue, top; L4: purple, upper middle; L5/6: green, lower middle). Differences in time are shown at the bottom for each compartment at two alpha levels (filled: 0.05; unfilled: 0.1) as computed by a two-sample t test. Average information transmission during the time of N2pc indicated with bars at right with upper limit of 95% confidence intervals (left bars, selective columns; right bars, non-selective columns). Significance is indicated with a magenta bracket and p value from a two-sample t test shown to the right of ordinate.

To identify columns with significant selectivity for either red or green, we performed Wilcoxon sign rank tests between the distribution of ratios in each column against bootstrapped null distributions. Each bootstrapped null distribution contained 15 randomly selected ratios from the full dataset (450 experimental values). 1000 such distributions were generated to be tested against. The bootstrapped distributions represent the range of possible values observed across V4, but do not capture any difference in the homogeneity of feature selectivity within a column.

We found that more than half of V4 columns (monkey Ca: 12/21, 57.1%; He: 5/9, 55.6%) show color selectivity defined in this way. We computed the information transmission of target position for each of these color tuned columns. We first computed this value for trials where the preferred color was in the column’s population receptive field. Then, we recomputed this value for trials with the non-preferred color. Note that the amplitude of the extracranial signal during the N2pc window did not differ across sessions with different target and distractor colors (paired sample t(16) = 0.40, p = 0.69) (Figure 4C), nor did the laminar CSD during the same window (L2/3: t(16) = −0.85, p = 0.41; L4: t(16) = 0.75, p = 0.46; L5/6: t(16) = 0.36, p = 0.72) (Figure 4D). However, we found that the information transmitted during the N2pc window was greater when a preferred rather than a nonpreferred color was present (Figure 4G). This difference was significant in L2/3 and L5/6 but not in L4 (t test across time with at least 10 ms having p < 0.05), and can clearly be seen at the single session level (Figure S4).

We investigated whether the degree of color preference was related to information transmission. We plotted the columnar color preference as an index (CCSI: positive, red-preferring; negative, green-preferring) against the difference in information transmission between conditions (red – green) around the time of peak information transmission (160-180 ms) for each session (Figure 4H). Computing Spearman’s r for each laminar compartment, we found a significant relationship between the magnitude of feature selectivity and the difference in information transmission for L2/3 (R = 0.50, p = 0.005) and L5/6 (R = 0.51, p = 0.004).

In a similar vein, we tested whether feature selective columns, on average, transmitted more information than their non-feature-selective counterparts. We found that feature selective columns, along all laminar compartments, transmitted significantly more information (Figure 4I) (two-sample t test: L2/3, p = 0.044; L4, p = 0.023; L5/6, p = 0.009). Together these findings suggest that visual cortical columns contribute more to the overlying attention-associated electric field when the item in their receptive field matches their tuning preference.

### Translaminar currents in V4 recapitulate the N2pc

CSD is computed by differentiating between local field potentials to eliminate volume conducted signals that do not arise from local circuit activity. Using the inverse procedure (i.e., summing the CSD), it is possible to estimate the local field potential without contamination by volume conducted activity (Nicholson and Llinas, 1971; Kajikawa and Schroeder, 2011). We used this logic to compute an estimated extracranial event-related potential (ERP). Specifically, we computed the sum of currents produced by a cortical column to estimate the extracranial signal at a position directly above. The resultant potential (ERP_cal_) showed a significant difference that persisted throughout the time period of the N2pc (Figure 5). In other words, the summed potential generated by currents along V4 columns differentiates between attention conditions simultaneous with the extracranially measured attention-associated signal. Note that the shape of the of the empirically observed extracranial ERP (ERP_obs_) differs from the estimated extracranial ERP_cal_. This is expected in part because the ERP_obs_ reflects several more variables such as volume conducted contributions of nearby columns as well as the filtering and attenuating effects of the tissue and cranium above the gray matter (Nunez and Srinivasan, 2006). Given these expected differences, it is remarkable how well the difference in ERP_cal_ predicts the timing of the attention-associated electric field.

**Figure 5.**
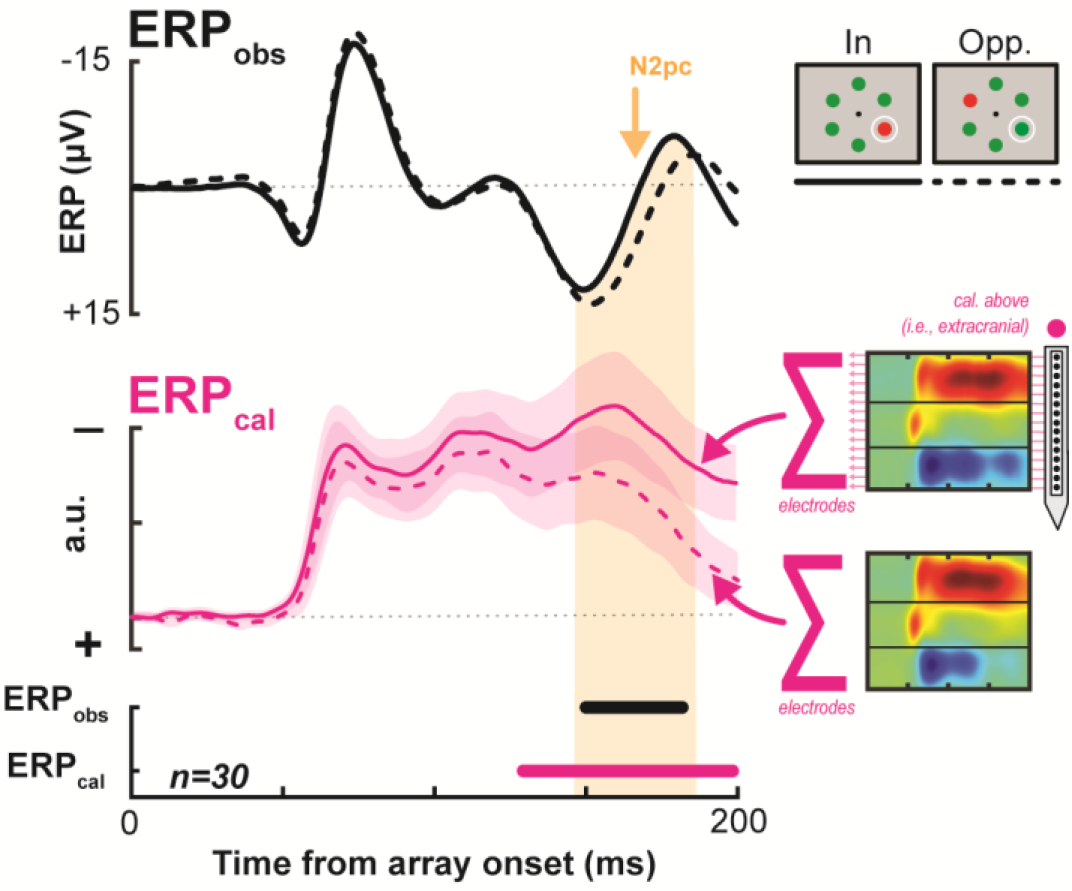
Comparing an estimated field potential generated from the CSD across the cortical columns to the actually observed extracranial event-related potential. Black lines indicate the empirically measured event-related potential (ERP_obs_, top), averaged across sessions. The pink line indicates the estimated event-related potential calculated from the synaptic currents across V4 columns, averaged across sessions (ERP_cal_, center). Synaptic currents at each electrode are measured and divided by the Euclidean distance of the electrode from the extracranial surface (see Methods; Nicholson and Llinas 1971; Kajikawa and Schroeder 2011). ERP for target present in the RF vs. target opposite the RF is shown as solid and dashed lines, respectively (example array for each condition shown at top right). Clouds around ERP_cal_ lines indicate 95% confidence intervals across sessions for each condition. Note that despite differences in overall waveshape (which are likely due to the fact that V4 is not the only contributor to the attention-independent, visually evoked ERP), the timing of differences within signal types can be compared. The congruence in polarization of the difference in potentials is of similar note.

**Figure 6.**
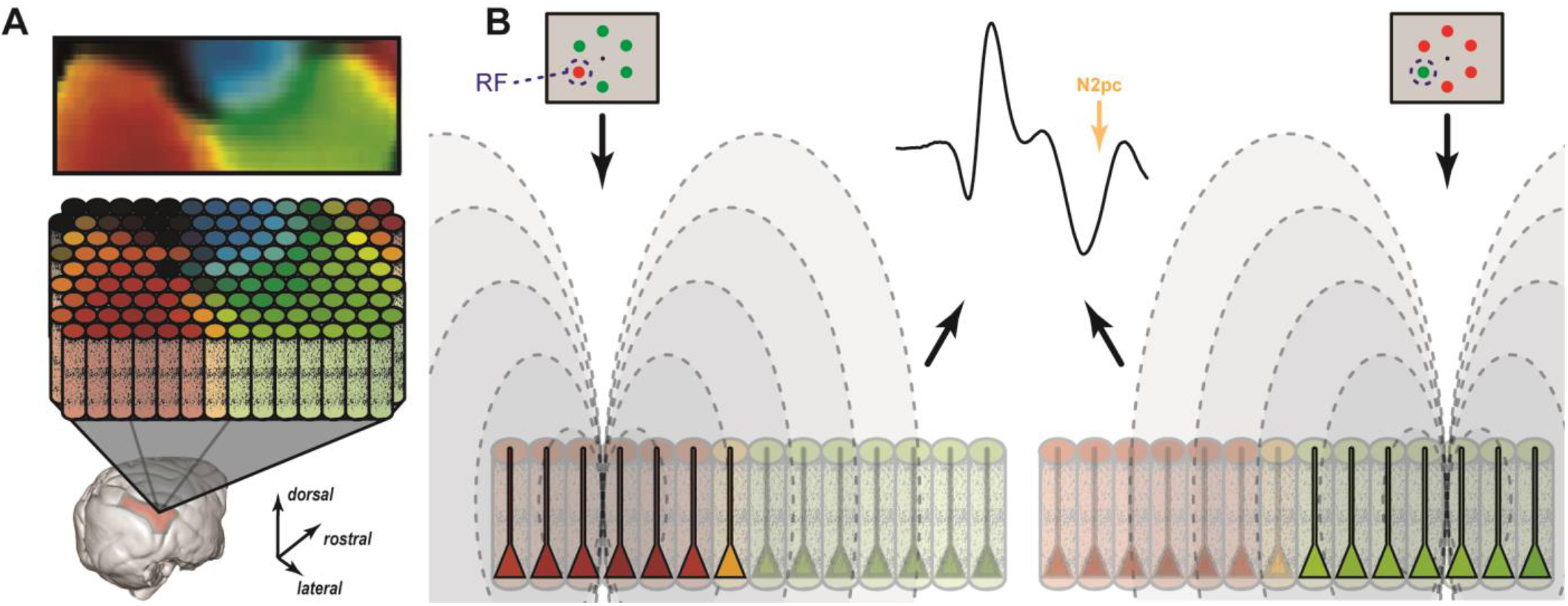
Feature Mosaic Hypothesis. (A) Top, a map of preferred color in area V4 derived from optical imaging (Tanigawa et al. 2010). Middle, a surface view map of columns extending through the layers of area V4. Bottom, this columnar structure was observed when n eural signals were sampled across all layers of area V4 on the surface of the prelunate gyrus highlighted red on structural brain scan obtained from monkey Z. (B) Relative contributions of cortical columns in area V4 to the attention-associated electric field when a red (left) or green (right) target appears in the RF. Intensity of pyramidal neuron activity is indicated by saturation in the diagram. The mesoscopic columns produce electric fields (dashed lines) that sum to produce the equivalent ERP.

## Discussion

Bioelectric potentials have practical and clinical applications when their generators are known. For example, the electrocardiogram is useful in medicine because the physiological process associated with each phase of polarization is understood. Likewise, the electroretinogram is useful because the cell layers associated with each polarization are understood. In contrast, human ERP components indexing cognitive operations will have limited utility until their neural generators are known.

The ERP indices of attention such as the N2pc or Pd are commonly used to assess the deployment of attention in human participants, but can also be observed in macaque monkeys, enabling systematic concurrent EEG and intracranial neurophysiological recordings. Our objective was to identify the neural generator of the attention-associated electric fields that comprise ERPs like the N2pc. Using inverse modeling of cranial surface EEG and laminar resolved CSD in a cortical area, we demonstrate that translaminar synaptic currents in visual cortical area V4 contribute to the generation of attention-associated electric fields during visual search. The dipole resulting in this electric field stemmed from layer-specific interactions in extragranular (top-down recipient) cortical layers. Unexpectedly, we discovered that the contribution of a cortical column to the overlying electric field depended on whether the visual feature in the RF matched the selectivity of the column – an important consideration in the mechanism producing EEG potentials that may not be observable through the macroscopic EEG signal alone.

### Columnar mosaic underlying EEG

Our discovery that dipoles established by synaptic currents in visual cortical columns underlie the generation of the attention-associated electric fields is consistent with the observation that ERP components such as the N2pc are largest over the occipital lobe in humans (Luck and Hillyard, 1990, 1994) and macaques (Woodman et al., 2007) and with human MEG studies (Hopf et al., 2000). While the objective of this investigation was to identify interlaminar interactions producing these attention-associated electric fields, the approach also affords the opportunity to better understand the neural circuitry that produces it. For example, we sought to understand how visual processing characteristics, in the form of feature selectivity, of cortical columns contributes to the generation of the electric field. Visual features, like color used in this study, can be decoded from extracranial signals like EEG (Sandhaeger et al., 2019; Sutterer et al., 2021). Visual cortical area V4 contains a functional map of hue along its surface (Tanigawa et al., 2011) with individual columns comprising the map preferring the same color (Zeki, 1973, 1980; Tootell et al., 2004; Conway and Tsao, 2009; Kotake et al., 2009). We demonstrate that color-feature selectivity was consistent along cortical depth and discovered that the contribution of a column to the global electric field was greater when the feature in the RF was the preferred feature of the column. Specifically, columns that preferred green (or red) contributed more to the electric field when the item in the RF was green (or red) rather than red (or green).

The implications of this unexpected finding are illustrated in Figure 8A-B, which portrays how an attention-associated electric field like the N2pc can arise from different populations of cortical columns. Columns with receptive fields enclosing the target and also being selective for the particular features of the target establish stronger dipoles than do columns either representing different parts of visual space or other visual features. If target position or target feature change, then the columns contributing the strongest dipoles change accordingly. It is important to note that columns contribute regardless of the feature (provided the attended target is in the RF), there is simply a greater relative contribution when the attended item is a visual feature preferred by the columnar microcircuit. It is an open question whether these shifts in the voltage distribution are measurable on the human scalp due to smearing of the signals as they propagate through the skull and scalp (Nunez and Srinivasan, 2006). Additionally, we do not know if this observation generalizes to other cortical areas or other ERP components. However, the discovery has this general implication: A given ERP can arise from qualitatively different neural circuit configurations. This implication entails specific limits on the nature of mechanistic inferences available from ERP measures.

### Plausible N2pc localization

The attention-associated electric field measured in our task is most likely representative of the commonly measured N2pc component of the EEG ERP. Given our findings regarding the functional architecture comprising attention-associated electric fields, it is conceivable that the N2pc arises from multiple, anatomically distinct cortical areas. That is, given the ubiquity of columnar architecture in sensory cortex and the specificity of visual feature representations to different cortical areas, electric dipoles formed across visual cortical layers could come about across multiple visual cortical areas with the relative contribution of each depending on the feature being attended to. This realization could help reconcile conflicting interpretations of the cognitive states and operations that are supposed to be indexed by the N2pc (Eimer, 1996; Kiss et al., 2008; Pagano and Mazza, 2012; Foster et al., 2020). Moreover, contributions from areas other than V4 are plausible because previous neurophysiological studies in macaques demonstrate attentional selection signals during visual search in the temporal (e.g., Sato, 1988), parietal (e.g., Bisley and Mirpour, 2019), and frontal (e.g., Thompson et al. 2005; Zhou and Desimone 2011) lobes. Of particular note, neuroimaging studies in humans indicate a contribution to the N2pc from posterior parietal cortex (Hopf et al., 2000). In the same vein, FEF neurons locate the target among distractors as early as, or even before, the N2pc arises (Cohen et al., 2009; Purcell et al., 2013). Given the interconnectivity of FEF and V4 (Schall et al., 1995; Ungerleider et al., 2008; Gregorio et al. 2012; Ninomiya et al. 2012), the frontal lobe thus could be the functional origin of an attentional selection signal communicated to V4 and other posterior areas (Armstrong and Moore, 2007; Ekstrom et al., 2009; Gregoriou et al., 2009, 2012; Marshall et al., 2015; Popov et al., 2017), which in turn generate the N2pc (Westerberg and Schall, 2021) which would be observable as the attention-associated electric field demonstrated in our data.

## Methods

### Animal Care

Procedures were in accordance with National Institutes of Health Guidelines, Association for Assessment and Accreditation of Laboratory Animal Care Guide for the Care and Use of Laboratory Animals, and approved by the Vanderbilt Institutional Animal Care and Use Committee following United States Department of Agriculture and Public Health Services policies. Animals were socially housed. Animals were on a 12-hour light-dark cycle and all experimental procedures were conducted in the daytime. Each monkey received nutrient-rich, primate-specific food pellets twice a day. Fresh produce and other forms of environmental enrichment were given at least five times a week.

### Surgical Procedures

Two male macaque monkeys (*Macaca mulatta* monkey Z, 12.5 kg; *Macaca radiata* monkey P, 9 kg) were implanted with head posts and skull-embedded EEG arrays using previously described techniques (Woodman et al., 2007). One monkey (monkey P) was implanted with a subconjunctive eye coil. Two male macaque monkeys (*Macaca radiata*; monkey Ca, 7.5 kg; He, 7.3 kg) were implanted with head posts and MR compatible recording chambers with craniotomy over V4. Anesthetic induction was performed with ketamine (10 mg/kg). Monkeys were then catheterized and intubated. Surgeries were conducted aseptically with animals under O_2_, isoflurane (1-5%) anesthesia. EKG, temperature, and respiration were monitored. Postoperative antibiotics and analgesics were administered. Further detail is documented elsewhere (Woodman et al., 2007; Westerberg et al., 2020a, 2020b).

### Magnetic Resonance Imaging

Anesthetized animals were placed in a 3 Tesla Magnetic Resonance Imaging (MRI) scanner. T1-weighted 3D MPRAGE scans were acquired with a 32-channel head coil equipped for SENSE imaging. Images were acquired using 0.5 mm isotropic voxel resolution with parameters: repetition 5 s, echo 2.5 ms, flip angle 7°.

### Visual Search Tasks

Monkeys performed a color pop-out (monkeys Ca, He, and Z) or T/L (monkey P) search. Search arrays were presented on a CRT monitor at 60 Hz, at 57 cm distance. Stimulus generation and timing were done with TEMPO (Reflective Computing). Event times were assessed with a photodiode on the CRT. We used isoluminant red and green disks on a gray background (pop-out) or uniform gray T’s and L’s on a black background (T/L). Target feature varied within session for monkeys Ca, He, and Z. Monkey P identified the same target on any given session (T or L) but changed specific targets session to session. Trials were initiated by fixating within 1 (monkeys Ca and He) or 2 (monkeys P and Z) degrees of visual angle (dva) of a fixation dot. Time between fixation and array onset was at least 500 ms (monkey P: 500–1000 ms; Z: 500 ms; Ca and He: 750–1250 ms). For monkeys experiencing a range of fixation periods (monkeys Ca, He, Z), a nonaging foreperiod function was used to determine the fixation period on a trial-by-trial basis. Arrays comprised of 6 (monkeys Ca and He) or 8 (monkeys P and Z) items. Monkeys P and Z experienced invariable array eccentricity (10 dva) and item size (monkey P: 1.3×1.3 dva; Z: 1×1 dva). 2 items were positioned on the vertical meridian, 2 on the horizontal, and the 4 remaining items equally spaced between. Monkeys Ca and He viewed items where size scaled with eccentricity at 0.3 dva per 1 dva eccentricity so that they were smaller than the average V4 receptive field (RF) (Freeman and Simoncelli, 2011). The angular position of items relative to fixation varied session to session so that 1 item was positioned at the center of the RF. Items were equally spaced relative to each other and located at the same eccentricity. Each trial, 1 array item was different from the others. Monkeys saccaded to the oddball within 1 (monkeys Ca and He) or 2 s (monkeys P and Z) and maintained fixation within 2–5 dva of the target for more than 400 ms (monkeys Ca, He, and Z: 500 ms; monkey P: 400–800 ms). Juice reward was administered following successfully completion of the trial. The target item had an equal probability of being located at any of the 6 or 8 locations. Eye movements were monitored at 1 kHz or 250 Hz using a corneal reflection system (monkeys Ca, He, and Z) or a scleral search coil (monkey P), respectively. If the monkey failed to saccade to the target, they experienced a timeout (1–5 s).

### 10-20 EEG Recordings

Two monkeys were implanted with an array of electrodes approximating the human 10-20 system locations (monkey P: FpFz, C3, C4, P3, P4, OL, OR, Oz; monkey Z: FpFz, Fpz, F3, F4, FCz, Cz, C3, C4, Pz, P5, P6, POz, O1, O2, Oz) (Jasper 1958). Referencing was done using either the FpFz electrode (monkey P) or through linked ears (Z). The impedance of the individual electrodes was confirmed to be between 2–5 kOhm at 30 Hz, resembling electrodes used for human EEG. EEG was recorded using a Multichannel Acquisition Processor (Plexon) at 1 kHz and filtered between 0.7–170 Hz. Data was aligned to array onset and baseline corrected by subtracting the average activity during the 50 ms preceding the array onset from all timepoints. Data was clipped 20 ms prior to saccade to eliminate eye movement artifacts.

### Simultaneous V4 CSD and Extracranial Recordings

The extracranial electric fields and laminar V4 neurophysiology were acquired at 24 kHz using a PZ5 and RZ2 (Tucker-Davis). Signals were filtered between 0.1-12 kHz. V4 data was collected from 2 monkeys (monkey Ca: left hemisphere; He: right) across 30 sessions (monkey Ca: 21; monkey He: 9) using 32-channel linear electrode arrays with 0.1 mm interelectrode spacing (Plexon) introduced through the intact dura mater each session. Arrays spanned layers of V4 with a subset of electrode contacts deliberately left outside of cortex. The extracranial electric field was derived from the most superficial electrode outside the brain filtered between 1–100 Hz. CSD was computed from the raw signal by taking the second spatial derivative along electrodes (Nicholson and Freeman, 1975; Schroeder et al., 1998; Mehta et al., 2000; Westerberg et al., 2019) and converting voltage to current (Logothetis et al., 2007). We computed the CSD by taking the second spatial derivative of the LFP:

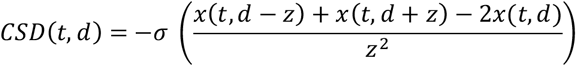

where *x* is the extracellular voltage at time *t* measured at an electrode contact at depth *d* and *z* is the inter-electrode distance and *σ* is conductivity. Both EEG and CSD were baseline corrected at the trial level by subtracting the average activation during the 300 ms preceding array onset from the response at all timepoints. Extracranial electric field potentials and CSD profiles were clipped 10 ms prior to saccade at the trial level to eliminate the influence of eye movements.

### Laminar Alignment

Orthogonal array penetrations were confirmed online through a reverse-correlation RF mapping procedure (Nandy et al., 2017; Westerberg et al., 2019; Cox et al., 2019a, 2019b; Dougherty et al., 2019) (Figure S6A). RFs were found to represent portions of visual space consistent with previous reports of V4 (Gattass et al., 1988) (Figure S6D). Positions of recording sites relative to V4 layers were determined using CSD (Schroeder et al., 1998; Nandy et al., 2017) (Figure S6B). Current sinks following visual stimulation first appear in the granular input layers of cortex, then ascend and descend to extragranular compartments. We computed CSD and identified the granular input sink session-wise. Sessions were aligned by this input sink (Figure S6C). ‘L4’ refers to granular input layer, ‘L2/3’ - supragranular layers, and ‘L5/6’ - infragranular layers. Each laminar compartment was assigned the same number of recording sites to alleviate biases during analysis.

### Inverse Modeling

Inverse modeling of 10-20 EEG recordings was performed in CURRY 8 (Compumedics Neuroscan). 3D head reconstruction was created for each monkey (P and Z) using the boundary element method (Hämäläinen and Sarvas, 1989). This method takes into account individual monkey’s surface morphologies to create models of cortex surface, inner and outer skull, and skin boundaries. This model was used in conjunction with EEG to compute a voltage distribution over the cortical surface. We calculated the current density with sLORETA, which calculates a minimum norm least squares that divides current by the size of its associated error bar, yielding F scores of activation. sLORETA produces blurred but accurate localizations of point sources (Pascaul-Marqui, 2002). Other algorithms such as Minimum Norm and SWARM were modeled as well, with agreement between models sufficient not to change any conclusions.

### Information Theory Analyses

Information theory (Shannon, 1948) analyses were chosen for several reasons. First, information theory analyses yield results in terms of ‘bits’ which can be used to directly compare effect sizes across measurement methods (e.g., CSD, Extracranial signal, and array composition [directed spatial attention]). Next, these analyses are inherently multivariate and able to capture linear and nonlinear relationships. Furthermore, information theory is model independent and does not necessitate a specific hypothetical structure in order to detect relationships between signals. This combination allows us to detect relationships between the extracranial signal and CSD signal that might not be linear and therefore would not be captured by linear models or correlation analyses. We chose to measure pairwise mutual information and information transmission to gauge the relationships between our three ‘signals’ (e.g., extracranial, CSD, and array composition [directed spatial attention]). Mutual information is the reduction in in uncertainty in one variable afforded by another known variable. That is, mutual information is greater when you know the state of one variable covaries with the state of the other variable. If the two variables do not correspond well, mutual information is low. Therefore, the reduction in uncertainty is formalized as ‘information’ which is relayed in bits. Mathematically, mutual information is captured by the following equation (Cover and Thomas, 2006; Beer and Williams, 2015):

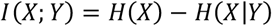

where *H*(*X*) and *H*(*X*|*Y*) are the entropy *X* and *X* given *Y*, respectively. Entropy for a signal (*S*) is computed by:

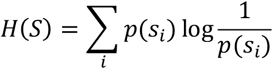

where *p*(*s*) is the probability distribution for signal *S* and *i* is the signal state. Therefore, mutual information can be computed probabilistically by:

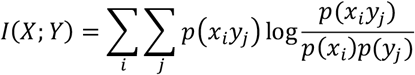

where *p*(*x*), *p*(*y*) are the probability distributions for *X* and *Y*, and *p*(*x, y*) is the joint probability distribution of *X* and *Y* across signal states *i* and *j* for *X* and *Y*, respectively.

While mutual information describes the relationship between the two signals (for our purpose: CSD and the extracranial signal, CSD and directed spatial attention, or the extracranial signal and directed spatial attention), it does not allow for the evaluation of two signals regarding a third (e.g., CSD and the extracranial signal regarding directed spatial attention). For analyses where we want to understand information regarding the allocation of directed attention from the synaptic currents in V4 to the extracranial signal we use a modified equation rooted in the same entropy/mutual information principles. In computing information transmission, we are interested in the information about *X* (directed spatial attention), transferred from *Y* (CSD in V4) to *Z* (extracranial signal) formalized as:

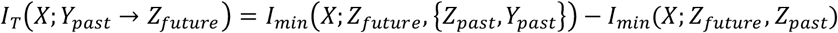

where *past* and *future* describe the timepoints when the data is taken from. The information transmission (*I*_*T*_) is taken as the difference between two minimum information calculations. The minimum information (*I*_*min*_) is computed regarding the combination of the individual signals (*S*_1_ and *S*_2_) at the specified time periods as:

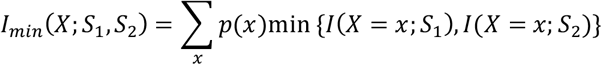

where *p*(*x*) is the probability distribution for signal *X* and *x* are the possible states of *X*. By taking into account different timepoints for the signals we can interpret this computation as the information about *X* (directed spatial attention) shared by *Y*_*past*_ (e.g., earlier CSD in V4) and *Z*_*future*_ (e.g., later extracranial signal) that was not already in *Z*_*past*_ (e.g., earlier extracranial signal).

Above information theory analyses were performed using the Neuroscience Information Theory Toolbox (Timme and Lapish, 2018). Pairwise mutual information and information transmission were computed at each timepoint across trials for each session using default parameters. Five uniform count bins were used for data binning. 10 ms was used for time lag for information transmission. Only correct trials were included. CSD for each laminar compartment was computed by taking the average activity of 5 sites at the trial level included in each laminar compartment. For mutual information between target position and the extracranial signal, target position was binary where target was either contra- or ipsilaterally presented. For computations within V4, target position was binary where target was either in the RF or positioned opposite the RF. 5000 Monte Carlo simulations were used to generate a distribution of null model values which experimental values were compared to (α = 0.05).

### Feature Selectivity

For each recording site within a column, gamma power (30-90 Hz) (Maier et al., 2010) responses were computed when either a red distractor was presented to the RF of the column or when a green distractor was present to the RF. Responses were taken as the average activation 75–200 ms following array onset. An index was computed from these responses by subtracting the two and dividing by their sum (CSI). Values were therefore bounded between −1 and 1 where larger magnitude indicates greater selectivity for green (towards −1) or red (towards 1). Columnar color selectivity index (CCSI) was computed as the average of CSIs along the entire column. We performed Wilcoxon signed rank tests on the distribution of CSIs across the recording sites of a given cortical column to determine whether a column was significantly color selective (α = 0.05). The selective columns were included in feature selectivity analyses where the preferred color and non-preferred color were defined as the color that elicited greater and lesser responses, respectively.

### Estimating Field Potential from CSD

We calculated the event-related potential at arbitrary positions from the measured laminar CSD (ERP_cal_) using a previously described model (Nicholson and Llinas, 1971; Kajikawa and Schroeder, 2011)

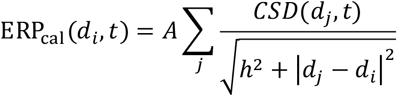

where ERP_cal_ at depth *i* (*d*_*i*_) for each timepoint (*t*) is taken as the sum of CSD at depths *j* (*d*_*j*_) for each timepoint divided by the Euclidean distance to account for the diminishing impact of local currents on more distant field potentials. The factor *A* acts only as a scaling factor and we cannot accurately estimate the magnitude of the one-dimensional CSD-derived waveform, so we eliminate this parameter from the calculation. This omission is consistent with previous reports (Kajikawa and Schroeder, 2011) and limits our comparisons of observed ERP and ERP_cal_ to only shape. However, magnitude differences can be observed between conditions for ERP_obs_ and ERP_cal_, independently. Also, for our purposes, we set *h* to 0 as we assume that our observed CSD and the calculated ERP are in the same vertical plane.

### Data Availability

Data supporting the findings documented in this study are freely available online through Dryad at https://doi.org/10.5061/dryad.djh9w0w15.

## Supplemental Information

Five additional supplementary figures are included to complement and expand primary findings.

## Acknowledgements

This work was supported by NIH through the NEI (P30EY008126, R01EY019882, R01EY008890, R01EY027402) and the Office of the Director (S10OD021771). J. A. W. was supported by fellowships from NEI (F31EY031293 and T32EY007135). The authors would like to thank I. Haniff, M. Feurtado, M. Maddox, S. Motorny, D. Richardson, L. Toy, B. Williams, R. Williams for technical support. The authors would like to thank B. Purcell., P. Weigand for collecting data and S. Errington, B. Herrera, K. Lowe, T. Reppert, J. Riera, A. Sajad, E. Sigworth for useful conversations regarding the work.

## Author Contributions

J. A. W., A. M., G. F. W., and J. D. S designed the research. J. A. W., M. S. S., and J. D. S. analyzed the data. J. A. W., M. S. S., and G. F. W. performed research. J. A. W. prepared visualizations. J. A. W., M. S. S., A. M., G. F. W., and J. D. S. wrote the manuscript.

## Supplementary Figures and Supplementary Figure Legends

**Figure S1.**
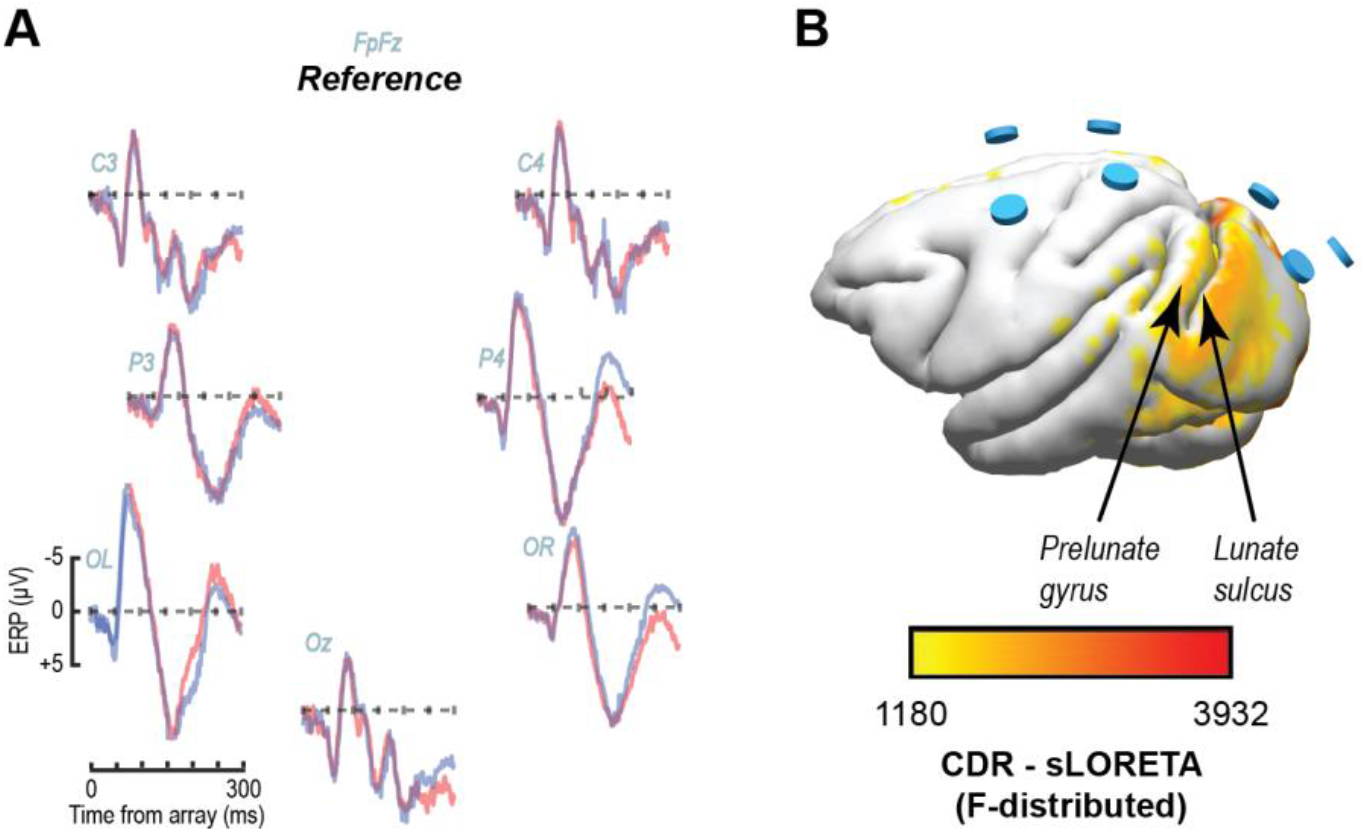
N2pc Distribution of monkey P (10-20 EEG recordings). (A) EEG traces for right (blue) and left (red) visual hemifield target presentations. Organization of traces reflects electrode positions. Scale is consistent across traces and is indicated by OL. N2pc was found to be significant through an ANOVA measured as the interaction between posterior electrode sites, the target position in the array, and the set size si tes (sites OR and OL, F(2,42) = 8.39, p < 0.001). (B) Inverse solution using sLORETA for N2pc (mean 190-300 ms following array onset) during a right visual hemifield target presentation displayed over the 3D render of MR scan for monkey P. Data clipped below 30% maximum value. Cyan cylinders indicate EEG electrode positions.

**Figure S2.**
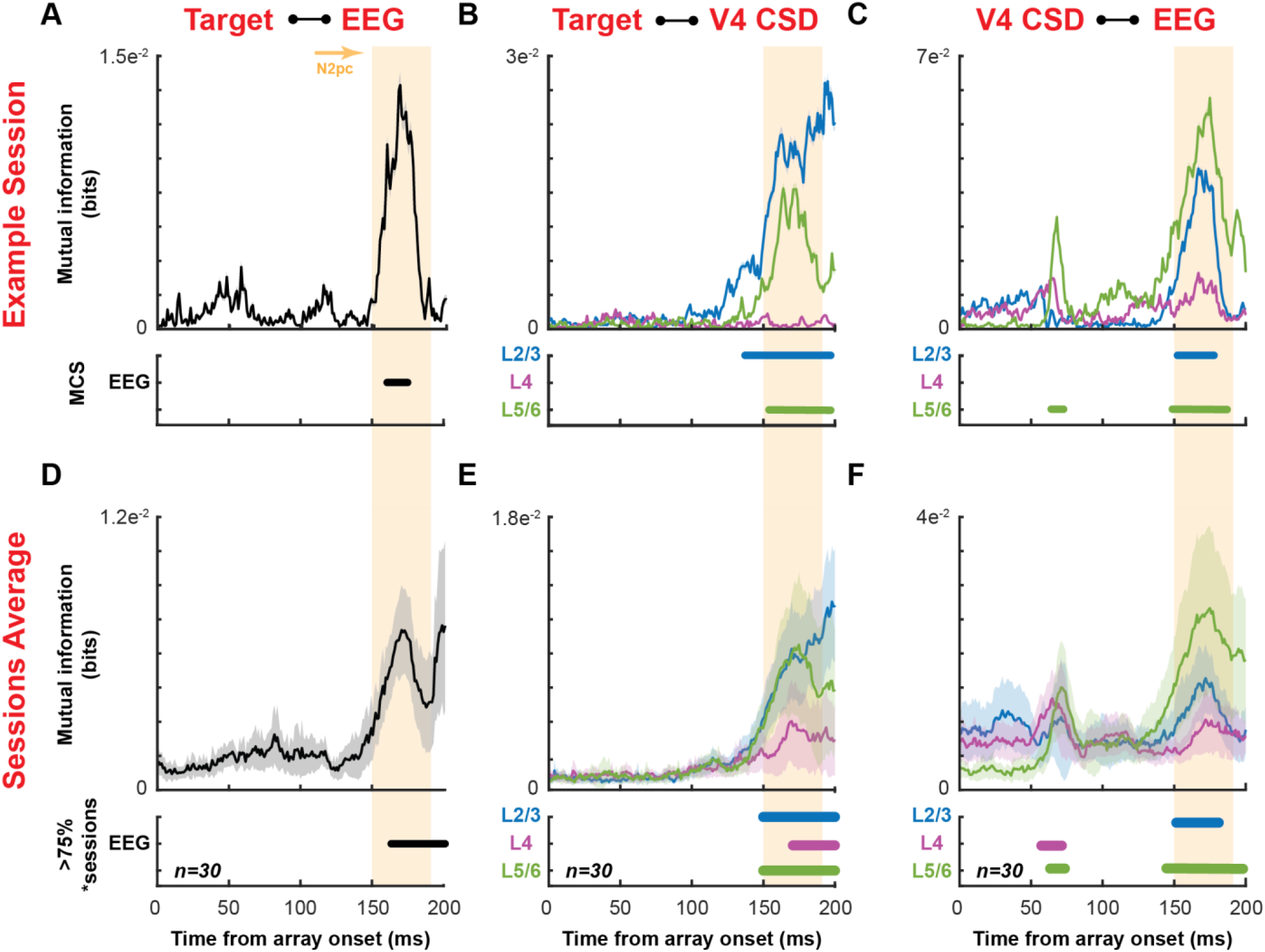
Mutual information measures for the extracranial signal, V4 CSD, and target position. (A) Mutual information between target position (binarily coded contra- or ipsi-presentation) and the extracranial signal along time (top) aligned on array onset with 95% CI cloud estimated from subsampling 75% of the data 100 times and recomputing. Significance established through Monte Carlo simulations is indicated below. Only epochs where significance >10 ms were included. Orange region indicates N2pc. (B) Mutual information between target position (binarily coded inside or opposite column RF) and each laminar compartment (L2/3 (blue), L4 (purple), and L5/6 (green)). Panel organization identical to (A). (C) Mutual information between the extracranial signal and each laminar compartment. Panel organization identical to (A). (D-F) Population averages (n=30) mutual information measures. Same organization as representative session, (A-C) respectively. Clouds around averages denote 95% confidence interval (CI) across sessions. Statistical measures for population averages reflect epoch’s where 75% sessions were found to be significant through Monte Carlo simulations for >10 ms.

**Figure S3.**
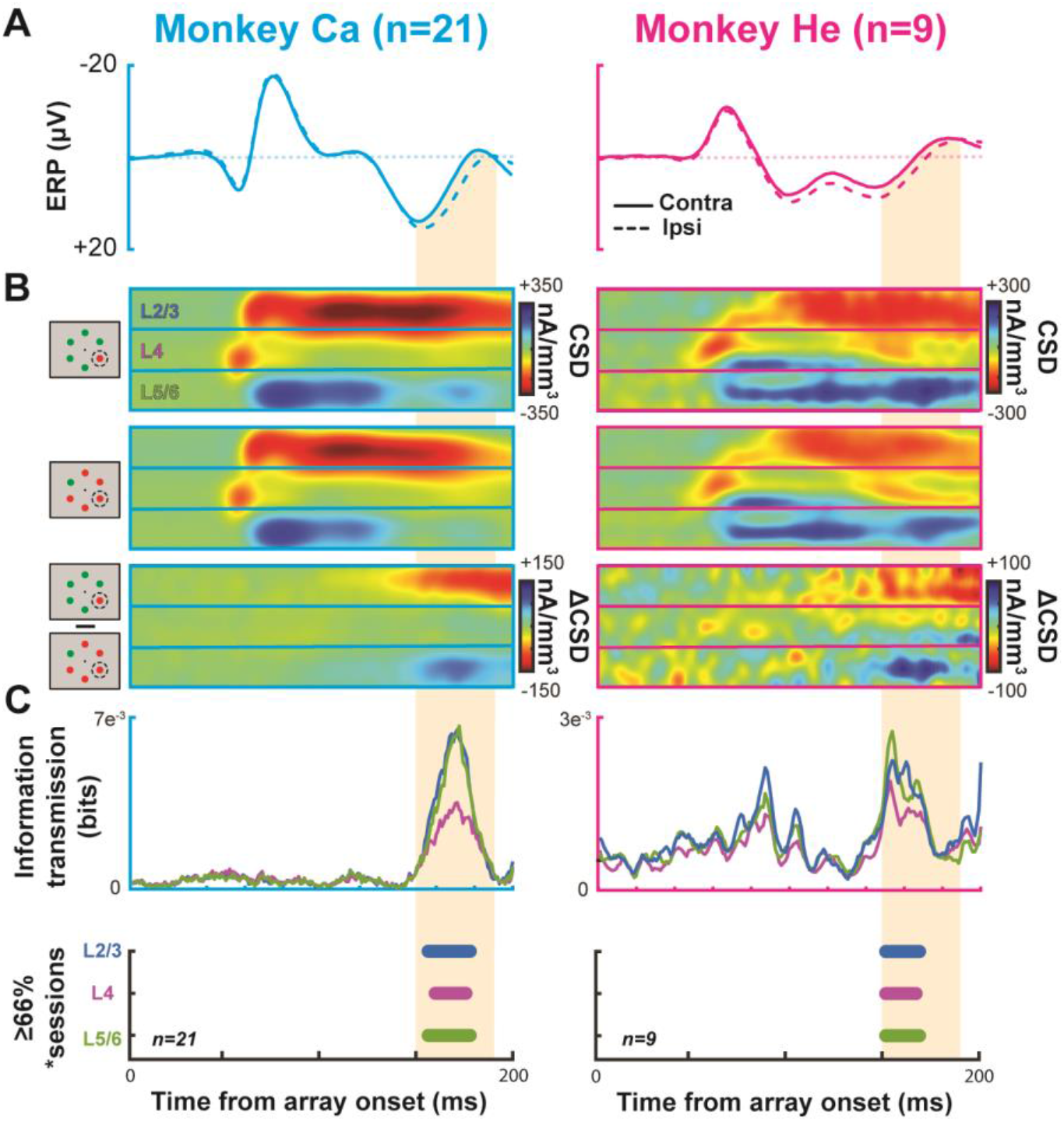
Individual monkey physiology and information transmission. Results for monkey Ca (n = 21) in left column and He (n = 9) in right column. (A) extracranial signal traces for target contralateral (solid line) and ipsilateral (dashed line) to recording site. Orange highlight represents the average time of N2pc used throughout the rest of the manuscript (150 – 190 ms). (B) Current source density profile for target in RF (top), outside RF (center), and the difference between the two (bottom). Horizontal boundaries indicate laminar compartments. (C) Information transmission regarding target position from laminar CSD to the extracranial signal (top). Blue represents L2/3, purple represents L4, and green represents L5/6. Timepoints where 66% of recorded sessions showed significant information transmission for more than 10 consecutive milliseconds through Monte Carlo simulations for each laminar compartment are shown at the bottom.

**Figure S4.**
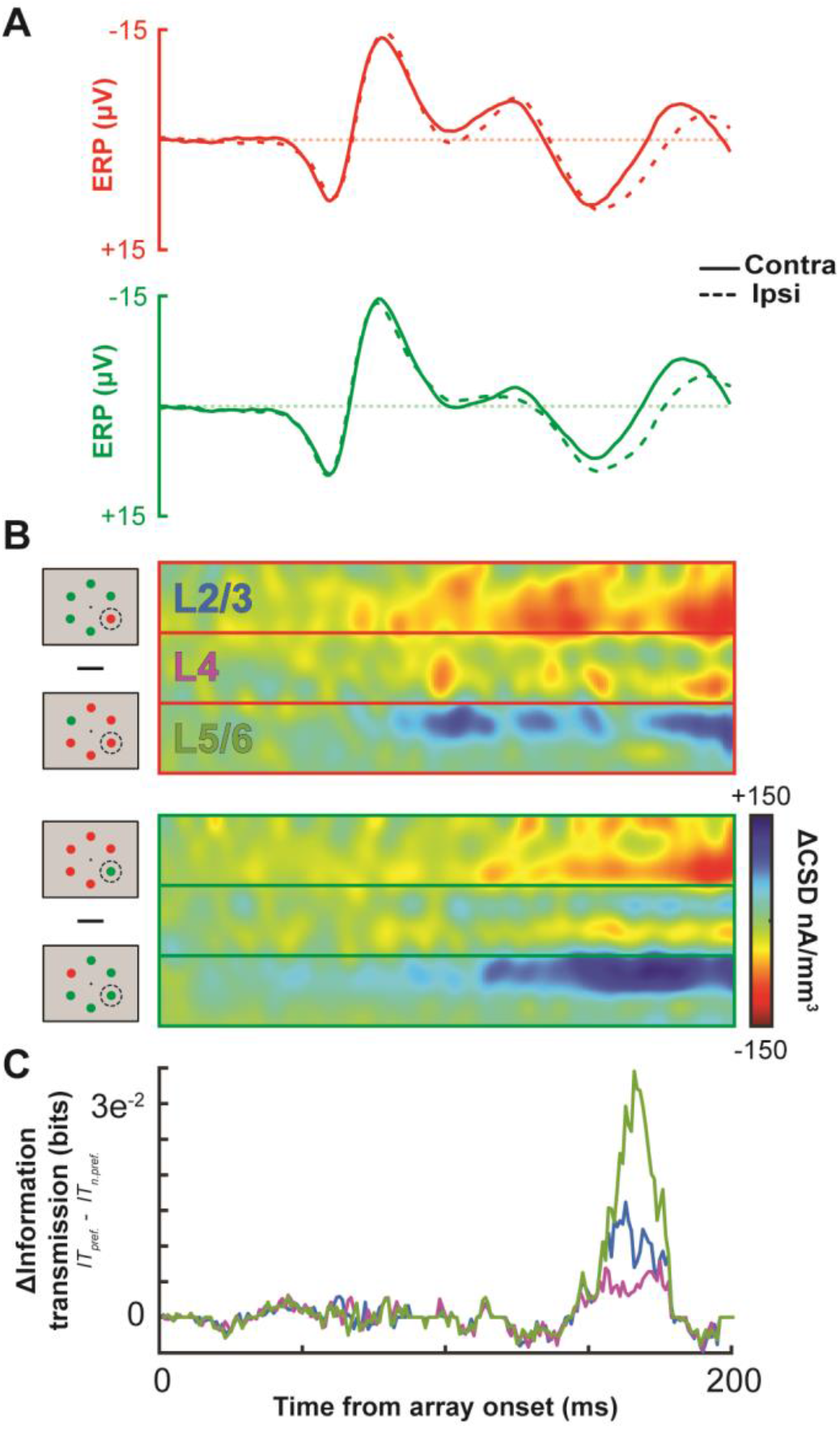
Single session example (monkey Ca) of the observed difference in information transmission depending on columnar color preference. (A) Extracranial event-related potential averaged across correctly performed trials (n=2992) for a single session with the target stimulus presented contralateral to the recording electrode (solid line; n_red_ = 742, n_green_ = 766) or ipsilateral to the recording electrode (dashed line; n_red_ = 752, n_green_ = 732) for trials where the target is red (top) of green (bottom). (B) Average difference in CSD profile for correctly performed trials between target present in RF and distractor present in RF for red item in RF trials (top) and green item in RF trials (bottom). (C) Difference in information transmission between the preferred color and the non-preferred color for the same single session as in panels A and B.

**Figure S5.**
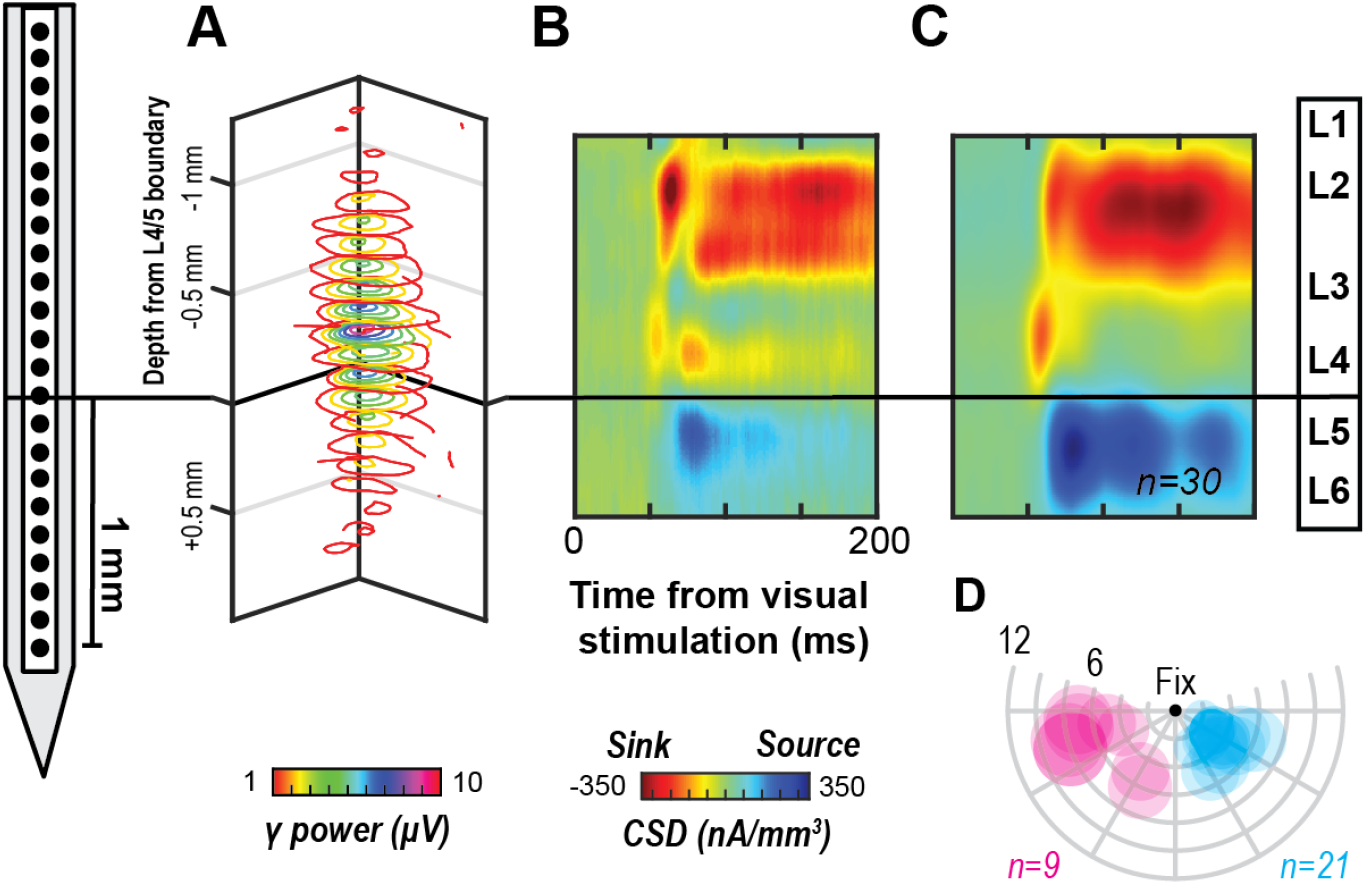
(A) Representative RFs across recording sites of a single array penetration. RFs across recording sites (z axis) are well aligned, indicating perpendicular penetration. Electrode positioned at left for reference. (B) CSD profile for the same session as (C). The initial sink following visual stimulation was used as a functional marker to determine the layer 4/5 boundary. Current sinks are indicated in red and sources in blue. The black horizontal line indicates the bottom of the granular input sink. Data are smoothed along depth and across time for visualization purposes. (C) Mean CSD profile following alignment of the 30 sessions (21, monkey C; 9, monkey H). Formatting identical to (B). (D) Columns’ RF locations across sessions and monkeys (cyan, monkey C; magenta, monkey H). RF centers determined online, and diameters estimated from previous reports (see V4 receptive field mapping and electrode orthogonality for details). Concentric circles indicate eccentricities in dva. Radial lines indicate angular positions relative to central fixation (black dot at top center).

